# Novel small RNAs expressed by *Bartonella bacilliformis* under multiple conditions reveal potential mechanisms for persistence in the sand fly vector and human host

**DOI:** 10.1101/2020.08.04.235903

**Authors:** Shaun Wachter, Linda D. Hicks, Rahul Raghavan, Michael F. Minnick

## Abstract

*Bartonella bacilliformis*, the etiological agent of Carrión’s disease, is a Gram-negative, facultative intracellular alphaproteobacterium. Carrión’s disease is an emerging but neglected tropical illness endemic to Peru, Colombia, and Ecuador. *B. bacilliformis* is spread between humans through the bite of female phlebotomine sand flies. As a result, the pathogen encounters significant and repeated environmental shifts during its life cycle, including changes in pH and temperature. In most bacteria, small non-coding RNAs (sRNAs) serve as effectors that may post-transcriptionally regulate the stress response to such changes. However, sRNAs have not been characterized in *B. bacilliformis*, to date. We therefore performed total RNA-sequencing analyses on *B. bacilliformis* grown *in vitro* then shifted to one of ten distinct conditions that simulate various environments encountered by the pathogen during its life cycle. From this, we identified 160 sRNAs significantly expressed under at least one of the conditions tested. sRNAs included the highly-conserved tmRNA, 6S RNA, RNase P RNA component, SRP RNA component, *ffH* leader RNA, and the alphaproteobacterial sRNAs αr45 and *speF* leader RNA. In addition, 153 other potential sRNAs of unknown function were discovered. Northern blot analysis was used to confirm the expression of eight novel sRNAs. We also characterized a <u>***B***</u>*artonella* <u>***b***</u>*acilliformis* **<u>g</u>**rou**<u>p I</u>** intron (BbgpI) that disrupts an un-annotated tRNA_CCU_^Arg^ gene and determined that the intron splices *in vivo* and self-splices *in vitro.* Furthermore, we demonstrated the molecular targeting of <u>***B***</u>*artonella* <u>***b***</u>*acilliformis* **<u>s</u>**mall **<u>R</u>**NA **<u>9</u>**(BbsR9) to transcripts of the *ftsH*, *nuoF*, and *gcvT* genes, *in vitro*.

**Author summary:** *B. bacilliformis* is a bacterial pathogen that is transmitted between humans by phlebotomine sand flies. Bacteria often express sRNAs to fine-tune the production of proteins involved in a wide array of biological processes. We cultured *B. bacilliformis in vitro* under standard conditions then shifted the pathogen for a period of time to ten distinct environments, including multiple temperatures, pH levels, and infections of human blood and human vascular endothelial cells. After RNA-sequencing, a manual transcriptome search identified 160 putative sRNAs, including seven highly-conserved sRNAs and 153 novel potential sRNAs. We then characterized two of the novel sRNAs, BbgpI and BbsR9. BbgpI is a group I intron (ribozyme) that self-splices and disrupts an unannotated gene coding for a transfer RNA (tRNA_CCU_^Arg^). BbsR9 is an intergenic sRNA expressed under conditions that simulate the sand fly. We found that BbsR9 targets transcripts of the *ftsH*, *nuoF*, and *gcvT* genes. Furthermore, we determined the specific sRNA-mRNA interactions responsible for BbsR9 binding to its target mRNAs through *in vitro* mutagenesis and binding assays.

## Introduction

*Bartonella bacilliformis* is a Gram-negative, facultative intracellular bacterium and the etiological agent of Carrión’s disease in humans. Carrión’s disease often manifests as a biphasic illness characterized by acute hemolytic anemia followed by eruptions of blood-filled hemangiomas of the skin [1]. Timely antibiotic administration restricts the fatality rate of Carrión’s disease to ~10%, although if left untreated, the rate has been reported to be as high as 88% [2, 3]. *B. bacilliformis* is transmitted between humans through the bite of female phlebotomine sand flies, specifically *Lutzomyia* spp. [4, 5]. The endemic region of Carrión’s disease has historically been limited to arid, high-altitude valleys (600 – 3200m) in the Andes Mountains of Peru, Colombia, and Ecuador, reflecting the habitat of the sand fly vector [6, 7].

The initial, acute stage of Carrión’s disease is referred to as Oroya fever (OF), and it is characterized by colonization of the entire circulatory system, leading to infection of ~61% of all circulating erythrocytes [7, 8]. This bacterial burden typically leads to severe anemia, fever, jaundice, and hepatomegaly, among other symptoms [9]. Weeks or months following OF, *B. bacilliformis* seemingly invades endothelial cells, where it triggers cell proliferation and angiogenesis. This event leads to formation of blood-filled blisters of the skin, referred to as verruga peruana (VP). The VP stage is chronic and lasts about one month to a year [1, 6]. Although Carrión’s disease can present as a severe illness, there are many documented cases with relatively milder symptoms and/or the onset of VP without having presented with OF symptoms [10]. In consideration of reports involving less virulent *B. bacilliformis* strains and the possibility that other *Bartonella* spp. can cause mild symptoms resembling Carrión’s disease, the incidence of the disease is likely underreported [11–13].

The *B. bacilliformis* infection cycle is strikingly under-studied compared to other vector-borne pathogens. It is clear that the bacterium is transmitted by *L. verrucarum* sand flies, although artificial feeding experiments showed that *L. longipalpis* can also “vector” the pathogen in the laboratory [5]. These studies also revealed that *B. bacilliformis* colonized and persisted in the lumen of the abdominal midgut of *L. verrucarum* but were digested along with the blood meal in *L. longipalpis* [5]. Despite this, viable bacteria were retrieved from both insects following a 7-d colonization period [5]. Other *Lutzomyia* spp. have been found to contain *B. bacilliformis* DNA, but colonization experiments have not been performed [14]. It has also been suggested that other mammals may serve as reservoir hosts for *B. bacilliformis*. However, serosurveys of animals that came into contact with infected humans were negative for *B. bacilliformis* DNA [15]. Interestingly, in various attempts to establish an animal model of *B. bacilliformis* infection, the bacterium was only able to infect rhesus macaques [16] and owl monkeys [17]. These results suggest that other primates could conceivably serve as natural reservoir hosts for *B. bacilliformis*, although there is a paucity of non-human primate species in *L. verrucarum*’s geographic range. Regardless, the lack of a small animal model severely limits the prospects of laboratory studies examining *B. bacilliformis* infections *in vivo*.

A number of virulence attributes are involved in *B. bacilliformis* pathogenesis, including erythrocyte attachment [18], invasion [19–21] and hemolysis [22]. Similarly, several factors have been implicated in endothelial cell invasion [23] and proliferation [24–26]. However, regulatory mechanisms that facilitate the pathogen’s virulence, colonization, and persistence in the sand fly have not been explored, to date. The disparate environments encountered by *B. bacilliformis* during transmission from sand fly vector to human host, and back again, suggest that genetic regulatory mechanisms are used to rapidly adapt to prevailing conditions. For example, the temperature of the sand fly vector would be comparable to ambient temperatures in the geographical range of the insect. The competent vector, *L. verrucarum*, is endemic to high-elevation ranges of the Occidental and Inter-Andean valleys of Peru, Colombia, and Ecuador [27], where temperatures range from 17°C - 22°C; fairly consistent with laboratory “room temperature” [28]. Upon transmission to the human host, the bacterium would need to adjust to a human body temperature of ~37°C. Similarly, human blood has a pH of ~7.4, while the pH of the sand fly (*L. longipalpis*) abdominal midgut after a blood meal is ~8.2, lowers to ~7.7 as the blood meal is digested, and decreases to ~6.0 after digestion [29, 30]. In contrast, the thoracic midgut is maintained at pH ~6.0, regardless of digestion status [29]. A rapid means of regulating virulence and stress-related factors to counteract sudden shifts in temperature and pH would be clearly adaptive for *B. bacilliformis*.

Bacteria often utilize small RNAs (sRNAs) to rapidly and efficiently regulate gene products involved in multiple biological processes. sRNAs are small (< 500 nts) non-coding transcripts that typically serve to up- or down-regulate translation of proteins by binding to the respective mRNA in a *cis* or *trans* fashion [reviewed in [31]]. This fine-tuning of protein production can enhance tolerance to stressors, including temperature [32] and pH [33]. To our knowledge, sRNA research in *Bartonella* is represented by a single report on *B. henselae* [34]. We therefore utilized total RNA-Sequencing (RNA-Seq) to interrogate *B. bacilliformis* transcriptomes to identify sRNAs expressed under a variety of conditions, including temperatures and pH levels consistent with the sand fly vector and human host. In doing so, we discovered 153 novel sRNAs expressed under at least one of the conditions tested. Furthermore, we characterized two of the sRNAs. The first RNA is a group I intron related to similar elements found in other alphaproteobacteria, while the other is a novel *Bartonella*-specific sRNA expressed only under conditions that simulate the sand fly vector.

## Materials and methods

### Bacterial culturing

Bacterial strains, primers, and plasmids utilized in this study are described in **S1 Table**. *B. bacilliformis* strain KC583 (passages #4-7) was cultivated on HIBB plates, comprised of Bacto heart infusion agar (Becton Dickinson; Franklin Lakes, NJ) containing 4% defibrinated sheep blood and 2% sheep serum (Quad Five, Ryegate, MT), by volume, as previously described [26]. Following 4 d of growth, 4 confluent *B. bacilliformis* plates per biological replicate were either shifted to different temperatures for 2 h, harvested and shifted to different pH levels in an HIBB liquid medium for 2 h, harvested and shifted to a human blood sample for 2 h, or harvested and used to infect cultured human umbilical vein endothelial cells (HUVECs; PCS-100-013; American Type Culture Collection; Manassas, VA) for 24 h. *Escherichia coli* (TOP10) was grown for 16 h at 37°C with shaking in lysogeny broth (LB), or on LB plates, supplemented with kanamycin (25 μg/ml) and ampicillin (100 μg/ml), when required.

### HUVEC culturing and infection

HUVECs were cultured and maintained as previously described [26]. *B. bacilliformis* infections were carried out for 24 h after which the medium was removed and cells were treated with gentamicin (10 μg/ml) for 1 h. Remaining viable extracellular *B. bacilliformis* cells were removed by washing 5 times for 10 min with phosphate-buffered saline (PBS; pH 7.4) solution. Finally, cells were harvested into TRI Reagent (Ambion; Austin, TX), as previously described for infected Vero cells [35].

### Human blood infection

Blood was drawn into vials containing sodium citrate to prevent coagulation. 1-ml aliquots were dispensed into fresh tubes, after which the lids were replaced with gas-permeable membranes. Blood samples were equilibrated at 37°C (HB37 samples) or 37°C in a blood-gas atmosphere (HBBG samples) for 1 h. Four HIBB plates of confluent *B. bacilliformis* for each equilibrated blood vial were harvested into PBS, pelleted at 16,000 x g for 5 min at 4°C and washed twice in PBS with identical centrifugation steps. Cell pellets were resuspended into 300 μl equilibrated blood, then dispensed back into the corresponding tube. The tubes were incubated at the appropriate condition for 2 h, then 1 ml RNALater solution (Thermo Fisher; Waltham, MA) was immediately added. Total RNA extraction was done as described below.

### Total RNA/genomic DNA isolation and preparation for RNA-Seq

Upon shifting *B. bacilliformis* for the designated time periods, cells were either harvested directly into one volume of RNAlater solution (Thermo Fisher) or centrifuged at 10,000 x g at room temperature for 2 min, after which the pellet was resuspended in a volume of RNAlater. The cells were incubated at room temperature for 1 h then frozen at −80°C for ≥ 2 h. The cells were thawed, centrifuged at 10,000 x g at 4°C for 10 min, and resuspended in 1 ml of TRI Reagent (Sigma-Aldrich; St. Louis, MO). The cells were incubated at room temperature for 1 h then frozen at −80°C for ≥ 2 h. Finally, cells were thawed and total RNA and genomic DNA isolation were done as previously described [35]. Total RNA pools from human blood infections were globin-depleted using a GLOBINclear kit (Ambion) according to manufacturer’s specifications. HUVE, HB37, and HBBG samples were enriched for bacterial RNA using a MICROBEnrich kit (Ambion). RNA (1 μg) from three independent biological replicates of each condition was sent to the Yale Center for Genomic Analysis (Pl25, Pl30, Pl37, pH06, pH07, and pH08 samples) or GENEWIZ (PlBG, HUVE, HB37, and HBBG samples) for bacterial rRNA depletion, stranded-library preparation, and HiSeq2500 (Illumina; San Diego, CA) 2×150 bp sequencing.

### Data analysis

Raw reads were quality filtered and aligned as previously described [35]. Briefly, raw fastq files were concatenated, quality filtered with the FASTX toolkit (http://hannonlab.cshl.edu/fastx_toolkit/), and then clipped, aligned, and filtered with Nesoni version 0.128 tools (http://www.vicbioinformatics.com/software.nesoni.shtml). Transcripts per million (TPM) were calculated using a custom Python script that can be accessed through GitHub (https://github.com/shawachter/TPM_Scripts). Stranded alignments were separated using SAMtools [36] and visualized using the Artemis genome browser [37].

sRNA identification was performed using the Artemis genome browser. RNA peaks were manually curated from IGRs and protein-coding gene regions. A read threshold for sRNA expression was devised for each condition tested based on reads that aligned to the *rpoD* gene (RpoD sigma factor), since TPM data suggest this gene is consistently expressed across all conditions. Using this method, putative sRNAs were identified, base ranges approximated, and sRNAs further characterized with putative promoters by manual searches using the conserved alphaproteobacterial sigma-70 promoter element, CTTGAC-N_17_-CTATAT [38]. Rho-independent terminators were identified using ARNold terminator prediction software (http://rna.igmors.u-psud.fr/toolbox/arnold/).

Since TPM calculations were done in the context of the total transcriptome and were not strand-specific, it was necessary to further refine TPM values for *cis*-anti sRNAs. This was done by considering the TPM value and proportion of the protein-coding gene to which the sRNA was antisense and subtracting the gene’s approximate TPM contribution with the following formula:

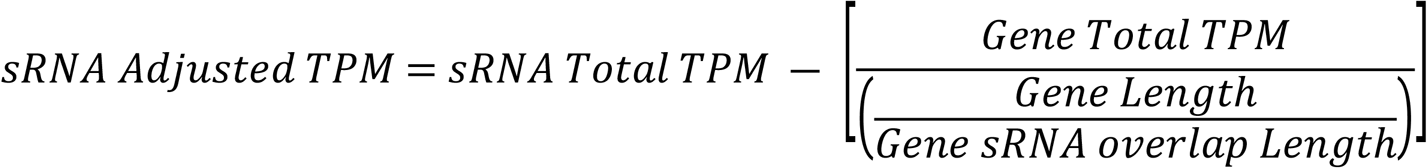

This was accomplished with a custom python script located in a GitHub repository (https://github.com/shawachter/TPM_Scripts). After this calculation, sRNAs whose TPMs were < 300 were considered not expressed for the purposes of the UpSet plot. This TPM value was chosen because it most accurately reflected the read threshold used in the initial manual sRNA search. All TPM values were included in the heatmap and determination of condition-specific sRNAs. Differentially-expressed sRNAs were determined by featureCounts [39] and the DESeq2 package of R version 3.4.4 [40]. For DESeq2 analysis, the p-value distribution of the significantly differentially expressed genes (DEGs) was re-sampled using fdrtools in order to more accurately achieve the desired null distribution [41]. This effectively made the analysis more stringent by providing fewer significant DEGs.

Infection-specific and sand fly-specific sRNAs were identified based on expression patterns as described in Results. The IntaRNA 2.0 sRNA target prediction algorithm was used to determine potential genes regulated by the sRNAs [42]. Only mRNA targets with a predicted IntaRNA 2.0 p-value < 0.01 were included in the potential targets list. Further targets with False Discovery Rate (FDR) values of < 0.05 were given special indications since these predicted bindings were considered especially strong. GO enrichment was performed utilizing the biobam Blast2GO program in the OmicsBox program suite (https://www.blast2go.com/blast2go-pro/download-b2g) using functional annotation of the *B. bacilliformis* KC583 genome as the background [43]. KEGG enrichment was performed using DAVID Bioinformatics Resources [44].

Figures were made using R version 3.4.4 and various Bioconductor packages including UpSetR [45], gplots (https://cran.r-project.org/web/packages/gplots/index.html), ggplot2 [46], and fdrtool [41]. Raw PNG images were modified into figures using Inkscape (https://inkscape.org/release/inkscape-0.92.4/) and Gimp (https://www.gimp.org/downloads/).

### Identification of transcription start sites

5’ RACE analyses of BbgpI and BbsR9 were performed using total RNA from *B. bacilliformis* shifted to liquid medium at pH 7 using a 5’ RACE System kit (Invitrogen; Carlsbad, CA) according to manufacturer’s protocols and with gene-specific primers (**S1 Table**). Resulting PCR products were cloned into pCR2.1-TOPO as instructed (Invitrogen), after which six arbitrary clones were sequenced with M13 universal primers by Sanger automated sequencing.

### Northern blots

Northern blot analyses were carried out using total RNA extracted from *B. bacilliformis* under the noted conditions. Northern blot probes were synthesized *in vitro* by engineering probe-specific PCR primers to contain a T7 promoter then utilizing a MAXIscript T7 Transcription kit (Invitrogen) supplemented with 0.5 mM Bio-16-UTP (Invitrogen). *B. bacilliformis* total RNA (2 μg) was resolved on a 1% denaturing agarose gel for 130 min at 57 V in 1X MOPS running buffer (Quality Biological; Gaithersburg, MD). The gel was washed in nuclease-free H_2_O for 10 min, followed by another wash in 20X SSC buffer (3M NaCl, 0.3M sodium citrate, pH 7.0) for 15 min. RNA was transferred overnight to a BrightStar-Plus nylon membrane (Ambion) in 20X SSC via upward capillary transfer. RNA was crosslinked to the membrane using a GS Gene Linker UV chamber (Bio-Rad; Hercules, CA) at 150 mJ. Membrane pre-hybridization and probe hybridization were done with a North2South Chemiluminescent Hybridization and Detection Kit (Thermo Fisher) according to manufacturer’s protocol. 50 ng of the appropriate *in vitro-* transcribed biotin-labeled probe was hybridized to the membrane at 67°C overnight. Membranes were washed 3 times for 15 min at 67°C in 1X Hybridization Stringency Wash Buffer (Thermo Fisher), developed, and imaged with a ChemiDoc XRS+ system (Bio-Rad).

### qRT-PCR

qRT-PCR was done on cDNA synthesized from 16 ng *B. bacilliformis* total RNA (for each 25 µl reaction) collected from various conditions using the Luna Universal One-Step RT-qPCR kit (New England BioLabs; Ipswich, MA) according to the manufacturer. *B. bacilliformis* total RNA was serially diluted and used as a standard curve, while primers targeting the *rpoD* housekeeping gene were used for normalization of gene expression between conditions. qRT-PCR was performed on a CFX Connect Real-Time System (Bio-Rad). cDNA from sRNAs of interest was analyzed for copy number, then divided by the copy number from the *rpoD* gene to achieve the sRNA transcripts / *rpoD* transcript values.

### Mutagenesis and RNA-RNA EMSAs

Mutagenesis of *gcvT*, *nuoF*, and *ftsH* target sequences was carried out *in vitro* using a Q5 mutagenesis kit (New England BioLabs) with specified primers (**S1 Table**). Primers engineered with a T7 promoter sequence were used to amplify the *gcvT* (−100 to +100), *nuoF* (−86 to +100), *ftsH* (−100 to +105), RS02100 (−76 to +100), *trmD* (−50 to +100), and *hflK* (−70 to +100) target sequences, where nucleotide +1 represents the first nucleotide of the protein-coding sequence. PCR products were cloned into pCR2.1-TOPO as instructed (Invitrogen). Resulting plasmid DNA was used as the template for Q5 mutagenesis. Q5 clones were sequenced, re-amplified with T7-engineered primers, and *in vitro* transcribed using the MAXIscript T7 Transcription kit (Invitrogen) with or without 0.5 mM Bio-16-UTP (Invitrogen), as required. Dose-dependent RNA-RNA EMSAs were performed as previously described [35] using 2 nM biotin-labeled BbsR9 and varying concentrations of *in vitro*-transcribed, unlabeled target RNA.

### Data availability

Aligned sequencing reads (BAM files) from all RNA-Seq experiments are available at the Sequencing Read Archive database (accession number PRJNA647605).

### Ethics statement

The Institutional Biosafety Committee at the University of Montana granted approval for the experimental use of human blood (IBC 2019-05). Formal consent was obtained in verbal form from the blood donor (co-author MFM).

## Results

### Identification of *B. bacilliformis* sRNAs

We first analyzed the total transcriptomic profiles of *B. bacilliformis* following a timed shift from normal culture conditions (4-d incubation on HIBB plates at 30°C) to various *in vitro* conditions that mimic the sand fly vector and human host (**Table 1**). Specifically, we controlled for several environmental variables, including temperature, pH, solid vs. liquid media, and the presence of a human blood-gas atmosphere. Following quality control analysis of the resulting RNA-Seq datasets and correlation of variation analysis (**S2 Table**), we discarded replicates that did not correlate well with others from the same condition. A principle component analysis (PCA) plot of the remaining RNA-Seq datasets confirmed statistical clustering of biological replicates (**S1 Fig**).

**Table 1.** Conditions used to prepare *B. bacilliformis* cultures for RNA-Seq experiments.

| Conditions | Medium | Designation | Shift Time | Simulation |
| --- | --- | --- | --- | --- |
| pH 7.4, 25°C | HIBB plates | PI25 | 2 hours | Sand fly ambient temperature |
| pH 7.4, 30°C | HIBB plates | PI30 | 2 hours | Sand fly ambient temperature |
| pH 7.4, 37°C | HIBB plates | PI37 | 2 hours | Human host |
| pH 7.4, 37°C with blood gas<sup>a</sup> | HIBB plates | PIBG | 2 hours | Human host |
| pH 6.0, 30°C | HIBB liquid | pH06 | 2 hours | Sand fly post-blood meal |
| pH 7.4, 30°C | HIBB liquid | pH07 | 2 hours | Human host / sand fly blood meal mid-digestion |
| pH 8.2, 30°C | HIBB liquid | pH08 | 2 hours | Sand fly initial blood meal |
| pH 7.4, 37°C with blood gas | HUVECs in EGM-Plus medium | HUVE | 24 hours | Human endothelial cell infection |
| pH 7.4, 37°C | Human blood | HB37 | 2 hours | Human erythrocyte infection |
| pH 7.4, 37°C with blood gas | Human blood | HBBG | 2 hours | Human erythrocyte infection |

HIBB, Bacto heart infusion blood agar containing 4% defibrinated sheep blood and 2% sheep serum (vol/vol); HUVECs, human umbilical vein endothelial cells; EGM-Plus (Lonza), endothelial cell growth medium containing 2% fetal bovine serum and bovine brain extract.
<sup>a</sup> Blood gas is comprised of 5% CO<sub>2</sub>, 2.5% O<sub>2</sub>, and 92.5% N<sub>2</sub> at 100% humidity to simulate human blood.

Next, we visualized alignments for each RNA-Seq dataset and manually curated transcript peaks that could correspond to novel sRNAs. Peaks were found in intergenic regions (IGRs), antisense to annotated genes (*cis*-anti) or as leader RNAs in 5’ untranslated regions (UTRs) of annotated genes. The peaks were further refined based on proximity to neighboring peaks and a threshold read coverage based on expression of a housekeeping gene, *rpoD* (encoding sigma factor RpoD). We discovered 160 potential sRNAs, including seven highly-conserved sRNAs of other bacteria/alphaproteobacteria (**S3 Table**). Of the 153 other potential sRNAs, 81 were located antisense to annotated genes, 57 were encoded in IGRs, and the remaining 15 were potential leader RNAs. Leader RNAs were included in the study because further analysis would be needed to determine if they are true leader RNAs, stand-alone sRNAs, or perhaps both. We also identified putative promoter elements for each identified sRNA based on approximated transcriptional start sites (TSS’s). Next, we constructed an UpSet plot to visualize the numbers of significantly expressed sRNAs shared between various combinations of conditions (**Fig 1**) [47]. Results of this analysis suggested that, while 19 of the 160 identified sRNAs were expressed regardless of circumstance, the majority of sRNAs were expressed under specific conditions. Following this, we calculated transcripts per million (TPM) for each sRNA under all ten conditions in the context of the total transcriptomes (**S4 Table**). TPM is a normalized measure of gene expression, and although it is not always appropriate to compare TPM values across different RNA-Seq experiments [48], we constructed a heatmap to get a broad sense of sRNA expression patterns (**S2 Fig**). These results revealed three distinct clusters of conditions with similar sRNA expression patterns and allowed us to identify interesting sRNAs for further characterization.

**Fig 1.**
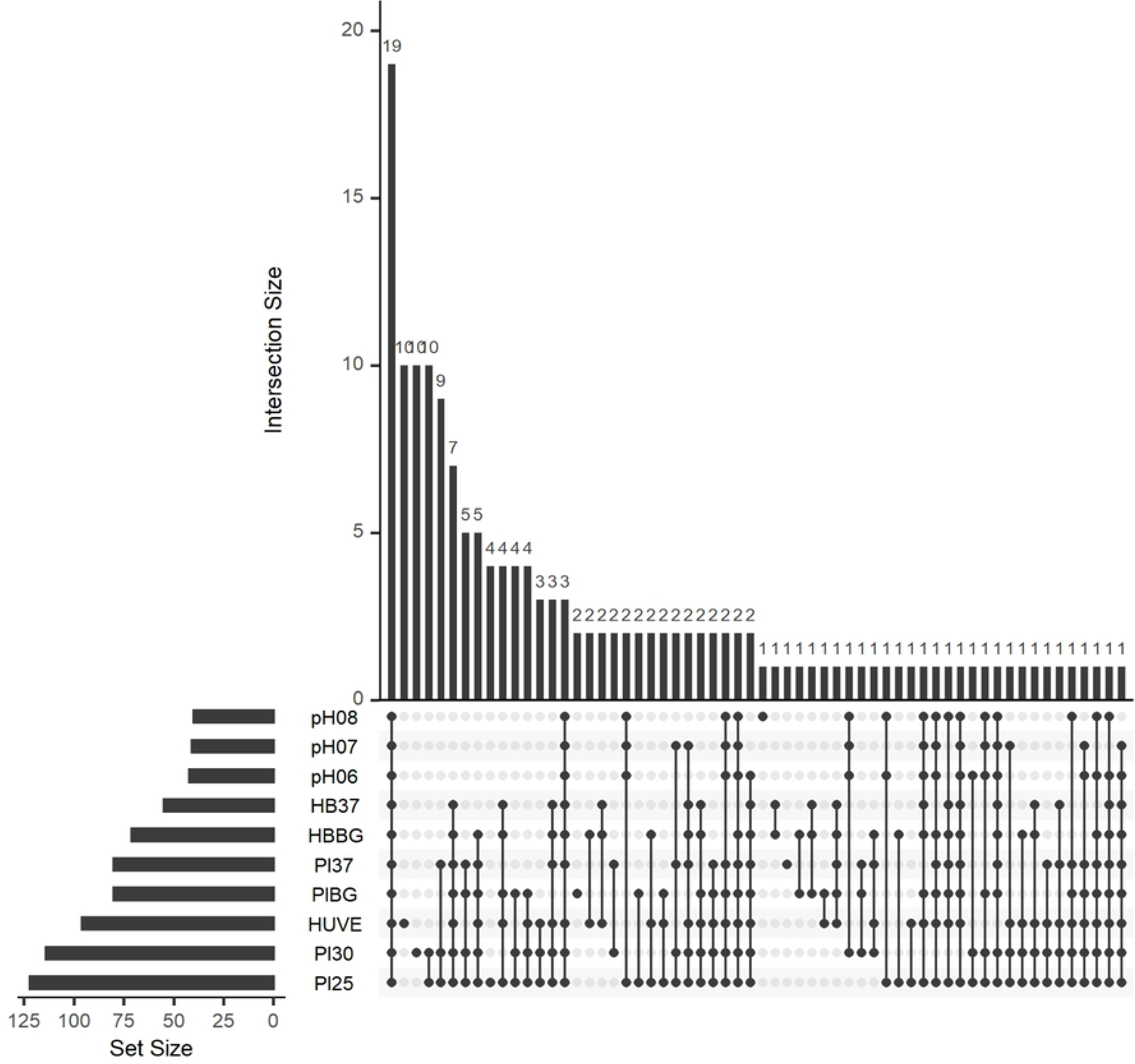
Most *B. bacilliformis* sRNAs are expressed under specific conditions. An UpSet plot is shown and displays the number of sRNAs shared among various combinations of conditions tested. The bar graph to the left indicates the quantity of sRNAs with a TPM >300 under the conditions shown. The connected nodes indicate shared conditions giving rise to the number of sRNAs expressed as indicated by the bar graph at the top.

### Verification of select *B. bacilliformis* sRNAs

Of the 160 putative sRNAs discovered in *B. bacilliformis*, we assigned name designations to 22 of them based on appraisal of general relevance, including six widely-conserved sRNAs, such as BbtmRNA (***<u>B</u>***. <u>***b***</u>*aciliformis* **<u>tmRNA</u>**), and 15 novel *Bartonella*-specific sRNAs. We also gave a name designation to the ***<u>B</u>***. <u>***b***</u>*aciliformis* **<u>g</u>**rou**<u>p I</u>** intron (BbgpI), a group I intron with related elements previously identified, but not characterized, in other alphaproteobacteria [49]. To corroborate RNA-Seq expression results and verify sRNA expression, we chose eight novel sRNAs at random and conducted Northern blot analyses. Results of the Northern blots confirmed the expression of all eight sRNAs (**Fig 2**).

**Fig 2.**
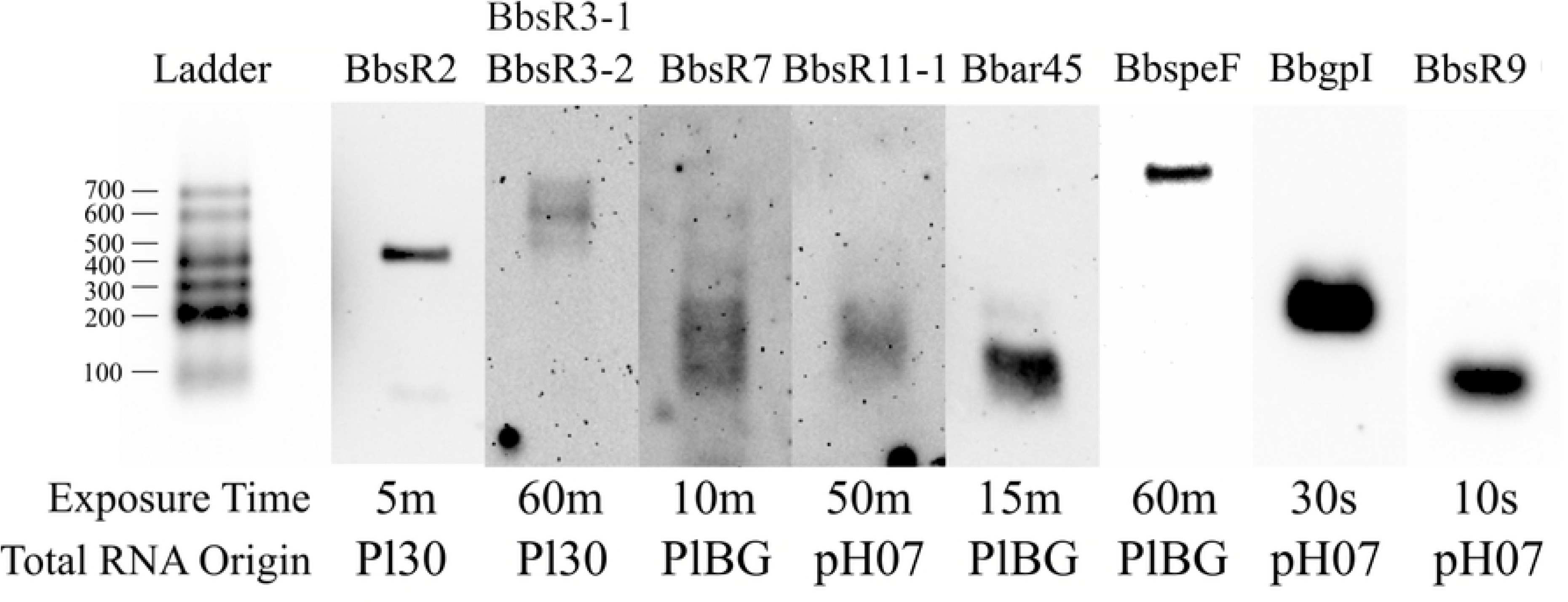
Northern blot analyses confirm expression of eight putative *B. bacilliformis* sRNAs. Eight separate Northern blots were run under identical experimental conditions (see Materials and methods). RNA ladders from the respective blots were aligned with each other for presentation of the resolved total RNA. sRNA designations are shown above each blot. Exposure times (minutes, m; seconds, s) and origin of the RNA are indicated below each blot.

We analyzed the differential expression of *B. bacilliformis* genes across the ten tested conditions by performing relevant pairwise comparisons (**Table 2**) using the DESeq2 package in R version 3.4.4 [40]. For this analysis, transcriptomes from all ten conditions were compared simultaneously, while specific, relevant pairwise comparisons were made (**Table 2**). Results showed the greatest number of significant differentially expressed sRNAs by comparing solid-to-liquid media and host cell types. Other sRNA candidates were also found to be differentially regulated by these comparisons (see **S3 Table**). We then utilized quantitative reverse transcription PCR (qRT-PCR) to validate the DESeq2 results. In doing so, we confirmed eleven sRNAs to be significantly differentially expressed under the relevant conditions (**S3 Fig**).

**Table 2.**
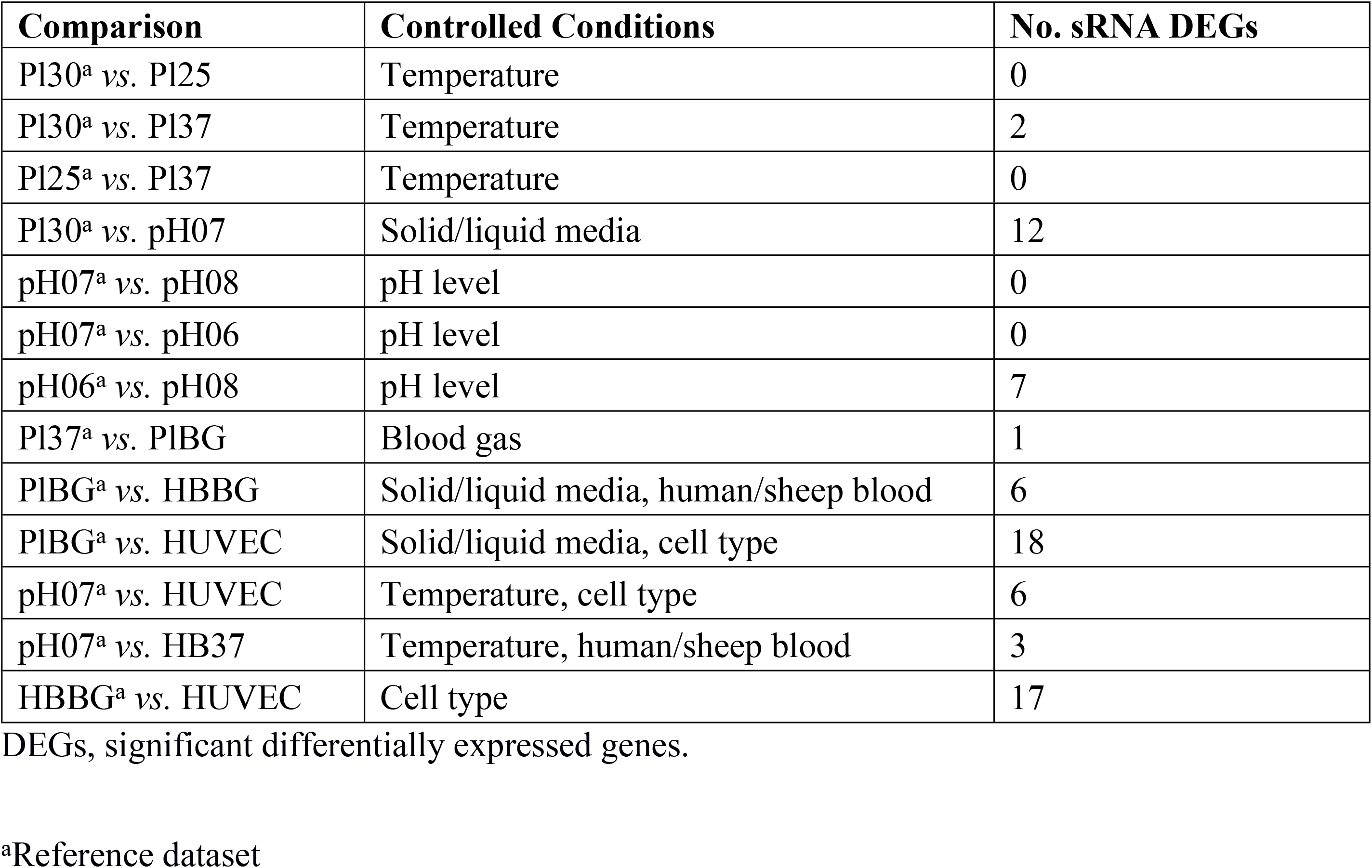
DESeq2 Comparisons Made.

### Condition-specific sRNAs target mRNAs enriched in specific pathways

We grouped several sRNAs based on their expression patterns across the ten conditions tested by using heatmap comparisons combined with data from the DESeq2 analysis. For example, multiple sRNAs were significantly and strictly expressed under conditions used to simulate or actually infect human cells (i.e., PlBG, HUVEC, HB37, and HBBG; see **Table 1**). These sRNAs were classified as human “infection-specific” based on their restricted upregulation (defined by TPM greater than the mean TPM plus one standard deviation) in at least two of these four conditions. Based on this definition, we identified 24 infection-specific sRNAs (see **S4 Table**).

We were also curious to determine if the predicted mRNA targets of the infection-specific sRNAs significantly corresponded to particular gene classifications to provide clues regarding their upregulation under infection-specific conditions. Using gene ontology (GO) and Kyoto Encyclopedia of Genes and Genomes (KEGG) enrichment analyses and the IntaRNA 2.0 sRNA target prediction program [42], we determined that the predicted mRNA targets of these sRNAs (**S5 Table**) were enriched for several GO terms, including protein/amide transport and nucleotidyltransferase activity (**Fig 3A**). The pool of mRNA targets was also enriched for the glycerophospholipid/glycerolipid metabolism and nucleotide excision repair KEGG pathways (**Fig 3B**).

**Fig 3.**
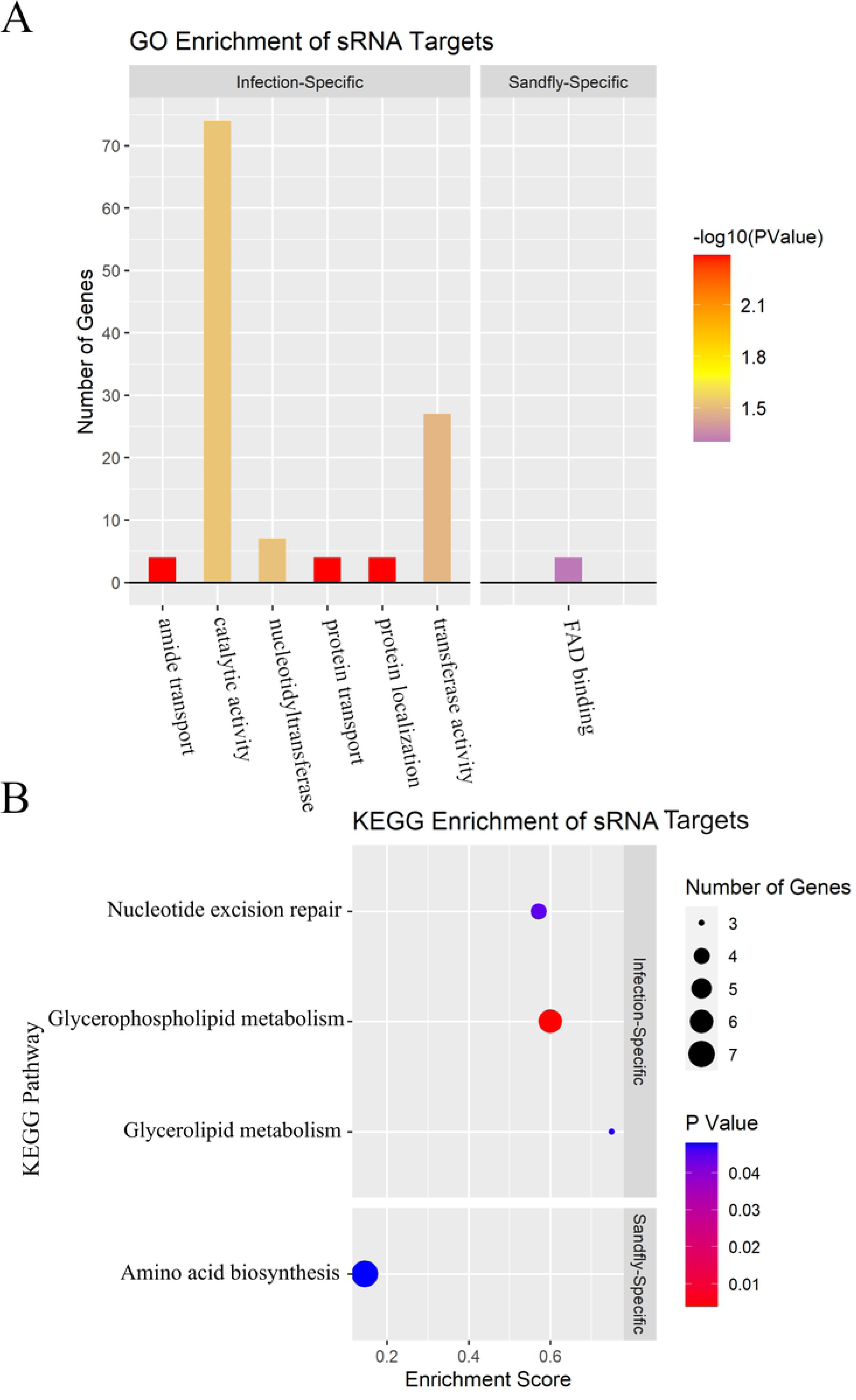
Condition-specific sRNA targets are enriched in several GO terms and KEGG pathways. **A)** Faceted bar graph of GO enrichment terms for infection-specific and sand fly-specific sRNA targets. Height of the bars indicates the number of sRNA targets containing that GO term, while the color displays the significance of enrichment. **B)** Faceted dot plot of KEGG enrichment terms for infection-specific and sand fly-specific sRNA targets. The enrichment score refers to the ratio of the number of gene targets corresponding to a particular pathway to the total number of genes in that pathway. Dot colors represent significance (p-value) of enrichment for that particular KEGG pathway.

We also determined that eight sRNAs were only expressed under conditions simulating the sand fly vector (i.e., Pl25, pH06, pH07, and pH08; see **Table 1**). Despite the fact that Pl25 clustered separately from pH06, pH07, and pH08 on the heatmap (**S2 Fig**), we included it in the sand fly-specific conditions due to upregulation of several sRNAs under both pH06/pH08 and Pl25 conditions (BB026-1, BB103-2, BB103-3, and BB124; **S4 Table**). These and other sRNAs were classified as “sand fly-specific sRNAs” based on their restricted upregulation in at least two of these four conditions. The predicted mRNA targets of the sand fly-specific sRNAs (**S6 Table**) were enriched for the flavin adenine dinucleotide (FAD) binding GO term (**Fig 3A**) and the amino acid biosynthesis KEGG pathway (**Fig 3B**).

### BbgpI is a group I intron that splices *in vivo* and self-splices *in vitro*

BB009 was initially identified as a sRNA of interest based on its high expression across multiple conditions (**S4 Table**). A BLAST search of the BB009 gene sequence (hereafter referred to as BbgpI) showed that it was highly homologous to a group I intron conserved in several other alphaproteobacteria and encoded in host tRNA_CCU_^Arg^ genes [49]. Since *B. bacilliformis* has no annotated tRNA_CCU_^Arg^ gene, we initially assumed that BbgpI was encoded in an IGR. However, such a location would be novel for group I introns, which are selfish genetic elements found in tRNA, rRNA, and rarely, protein-coding genes (reviewed in [50]).

To address this discrepancy, 5’ rapid amplification of cDNA ends (5’ RACE) was utilized to determine the 5’ end of the putative spliced-out RNA segment. From these results, we determined that BbgpI was flanked by CCT direct repeats, identical to those produced by the tRNA_CCU_^Arg^ alphaproteobacterial group I intron (**S4A Fig**) [49]. Also, the predicted secondary structure of BbgpI possessed conserved group I intron stem structures (**S4B Fig**). Finally, we scanned the locus and flanking sequences with the tRNAscan-SE 2.0 web server and identified a tRNA_CCU_^Arg^ gene (**S4C Fig**), but only when a sequence with BbgpI spliced out was used [51]. Taken together, these results suggest that BbgpI is a member of a conserved, alphaproteobacterial group I intron family and disrupts an unannotated tRNA_CCU_^Arg^ gene (locus: c42404-42711) of *B. bacilliformis*.

Although homology and structural results suggested that BbgpI was a group I intron, it was unclear whether BbgpI was able to self-splice or whether a protein cofactor was required [49]. To address this question, we examined BbgpI’s ribozyme activity *in vitro*. Following *in vitro* transcription, cDNA synthesis, and PCR analysis with the primers shown in **S4A Fig**, we determined that BbgpI self-splices *in vitro* and is spliced *in vivo* (**Fig 4**), in keeping with other group I introns.

**Fig 4.**
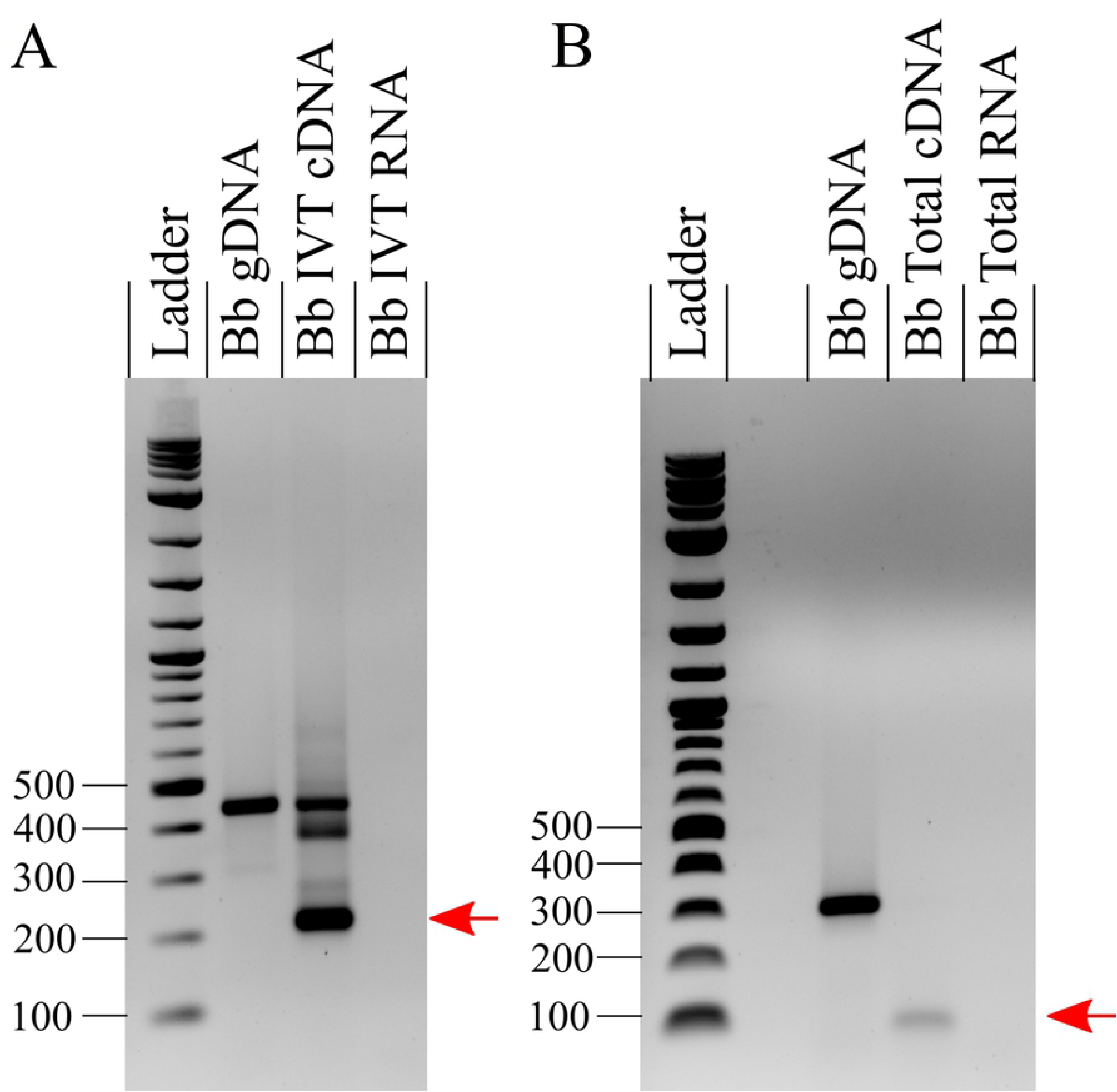
BbgpI self-splices *in vitro* and is spliced *in vivo*. **A)** PCR analysis of an *in vitro*-transcribed (IVT) region of *B. bacilliformis* (Bb) genomic DNA (gDNA) containing BbgpI. Ethidium bromide-stained agarose (1%) gels are shown. PCR on the resulting cDNA using “Nested Primers” (see **S4A Fig)** produced unspliced DNA (450-bp band), a partially-spliced product of ~390 bp, plus a 218-bp band corresponding to the BbgpI flanking region where BbgpI self-spliced out (indicated by the red arrow). PCR on a Bb IVT RNA negative control did not produce product. **B)** As in A) but utilizing cDNA synthesized from *B. bacilliformis* total RNA using “Splice Flank Primers” (see **S4A Fig)**. A 308-bp amplicon was produced from gDNA, whereas a 76-bp band, corresponding to the BbgpI flanking region with BbgpI spliced out, was produced from cDNA generated from total RNA (indicated by the red arrow). PCR on a Bb total RNA negative control did not produce product.

### BbsR9 is a sand fly-specific sRNA

BB092 (hereafter referred to as BbsR9) was initially identified as a sRNA of interest due to its restricted high-level expression under conditions that simulated the sand fly vector (**Table 1**, **S4 Table**). In addition, BbsR9 was found to have well-defined, predicted sigma-70 promoter and Rho-independent terminator regions, and it was conserved among several other *Bartonella* spp. (**S3 Table**). To elucidate the BbsR9 gene, the TSS was determined by 5’ RACE. These results showed two possible sites with equal representation among the six clones sequenced (**S5A Fig**). We therefore wished to determine if two distinct transcripts of BbsR9 were made or if there was a single, dominant transcript for the sRNA. In addition, we wanted to confirm BbsR9 expression across the conditions examined. To this end, we set out to determine if BbsR9 would remain highly expressed in sheep blood shifted to 37°C or if we would see a downregulation of the sRNA, as in *B. bacilliformis* shifted to human blood at 37°C (HB37/HBBG; see **S4 Table**). Northern blot analyses showed BbsR9 transcript and, together with the intensity of the signal, indicated a single, dominant TSS (bolded underlined in **S5A Fig**). Interestingly, we saw a distinct downregulation of BbsR9 when *B. bacilliformis* was shifted to sheep blood at 37°C compared to 30°C (**S5B Fig**). This decrease in RNA suggests that BbsR9 is primarily expressed under conditions that simulate the sand fly vector and not the human host. Taken as a whole, results of the Northern blots and RNA-Seq suggest that both a liquid medium (Pl30 vs. pH07; see **S4 Table**) and a temperature below 37°C upregulate BbsR9.

### BbsR9 targets transcripts of *ftsH*, *nuoF*, and *gcvT in vitro*

Since BbsR9 expression was restricted to sand fly-like conditions, we were interested in characterizing its mRNA targets to shed light on the sRNA’s role in regulation. To that end, we first utilized the TargetRNA2 [52], IntaRNA [42] and CopraRNA [53] algorithms to determine potential mRNA targets (**Table 3**). From these results, we selected transcripts of *ftsH*, *nuoF*, *gcvT*, *trmD*, *hflK*, and a predicted DNA response regulator (RS02100) as potential targets for characterization based on shared predictions between algorithms and the strength of predicted binding events.

**Table 3.**
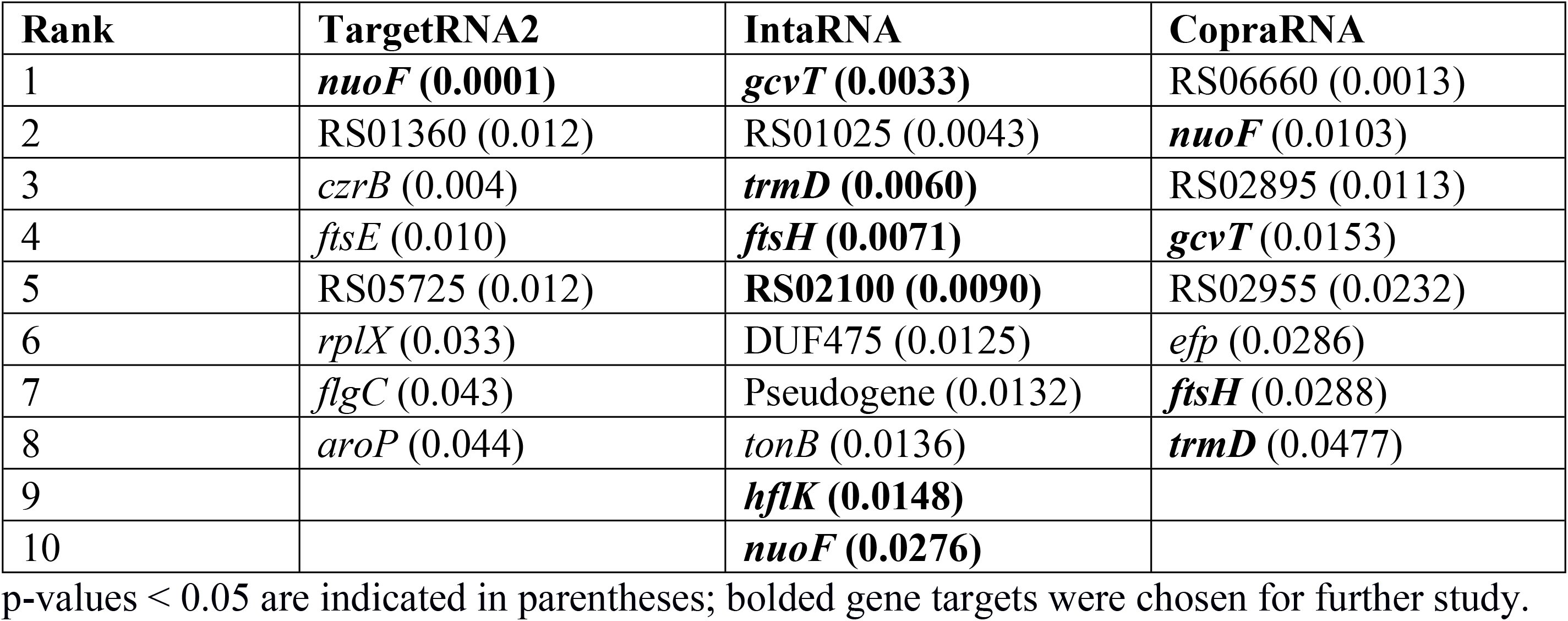
mRNA targets for BbsR9, as predicted by the indicated algorithms.

To demonstrate physical interactions between BbsR9 and the chosen mRNA candidates, RNA-RNA electrophoretic mobility shift assays (EMSAs) were done using *in vitro-*transcribed BbsR9 and segments of the target mRNAs of interest with their predicted sRNA target regions. Results of the EMSAs showed that BbsR9 bound mRNAs of *ftsH*, *nuoF*, and *gcvT in vitro*, as judged by the novel hybrid RNA species showing markedly slower migration during gel electrophoresis (**Fig 5**). Hybrid RNAs were not observed for the other three candidate mRNAs, suggesting that sRNA binding did not occur.

**Fig 5.**
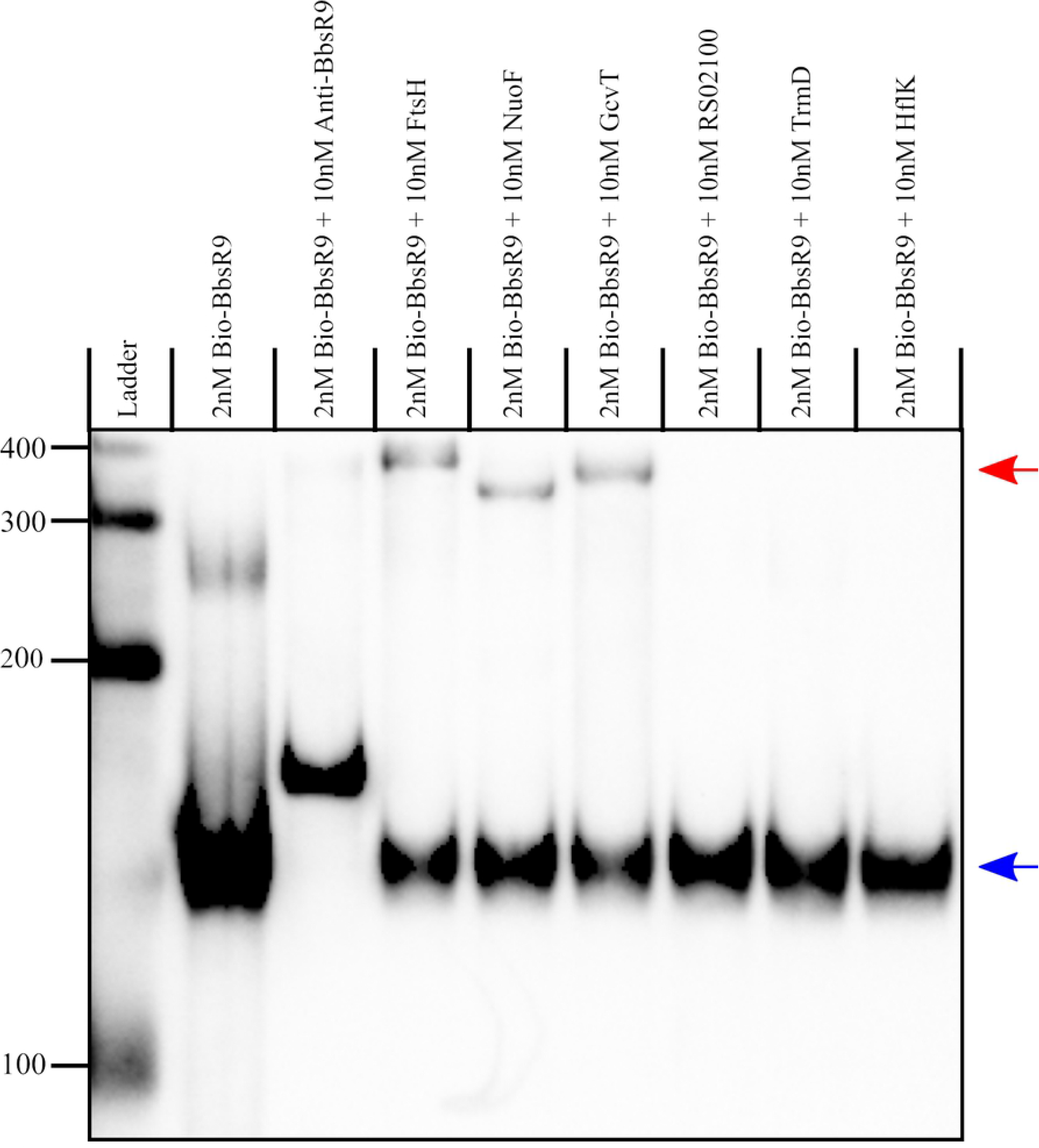
BbsR9 targets transcripts of *ftsH*, *nuoF* and *gcvT in vitro*. RNA-RNA EMSA of biotin-labeled *in vitro-*transcribed BbsR9 binding to *in vitro-*transcribed mRNA segments of the *ftsH*, *nuoF*, *gcvT*, *trmD*, BARBAKC583_RS02100 and *hflK* genes. Red and blue arrows indicate bands corresponding to BbsR9 bound and unbound to target RNAs, respectively. Base values of the RNA size standard (ladder) are shown on the left.

We further characterized BbsR9-mRNA interactions by mutagenizing the predicted sRNA-binding regions of *ftsH*, *nuoF*, and *gcvT* (**Fig 6**). The predicted *ftsH*-binding region was extensive, so we created two distinct mutants for this target as well as a double-mutant (**Fig 6**). RNA-RNA EMSAs conducted with the mutagenized target mRNAs showed complete elimination of BbsR9 binding to all three targets *in vitro* regardless of increasing target quantity present in the hybridization reaction (**Fig 7**). As expected, wild-type targets showed dose-dependent hybridization and signal intensity. Interestingly, abrogation of *ftsH* transcript binding by BbsR9 was only observed with mutation 1 (Mut1) alone or in combination with mutation 2 (Mut2), whereas Mut2 alone did not prevent BbsR9 binding to the RNA (**Fig 7**). In consideration of the RNA secondary structure predictions and the EMSA results, we conclude that BbsR9 primarily targets mRNA transcripts via multiple GC-rich regions of a large, predicted stem-loop structure (see **Fig 6**) [54].

**Fig 6.**
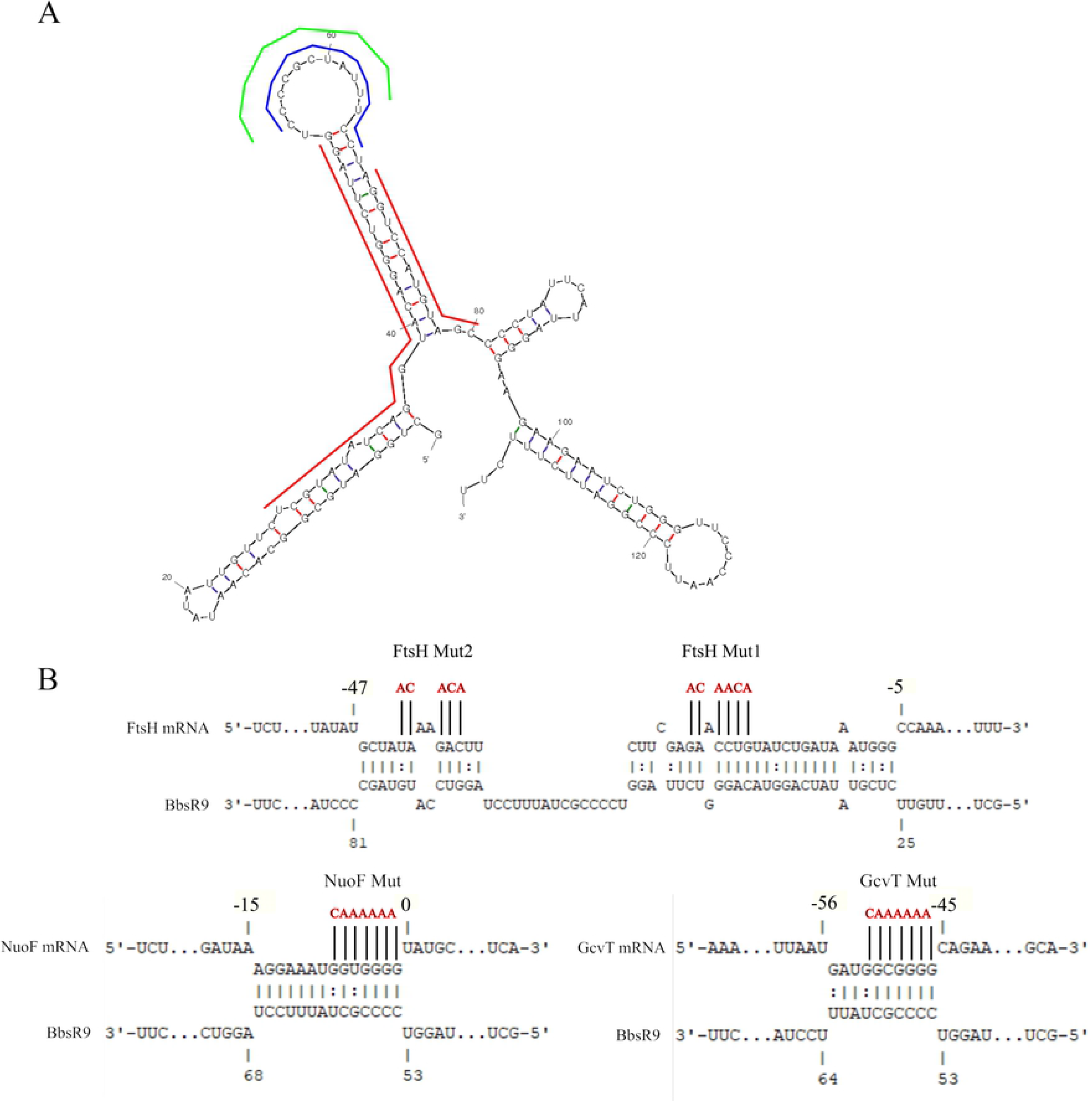
BbsR9 binds its targets through several GC-rich predicted seed regions. **A)** Mfold secondary structure prediction of BbsR9 (ΔG = −46.9 J mol-1) with predicted seed regions for *ftsH*, *nuoF*, and *gcvT* transcript binding indicated by red, blue, and green lines, respectively. **B)** Predicted IntaRNA BbsR9 target seed regions of the indicated transcripts. For the mRNA targets, nucleotide position +1 represents the first nucleotide of the respective start codon. Mutagenized bases of each mRNA are indicated in red.

**Fig 7.**
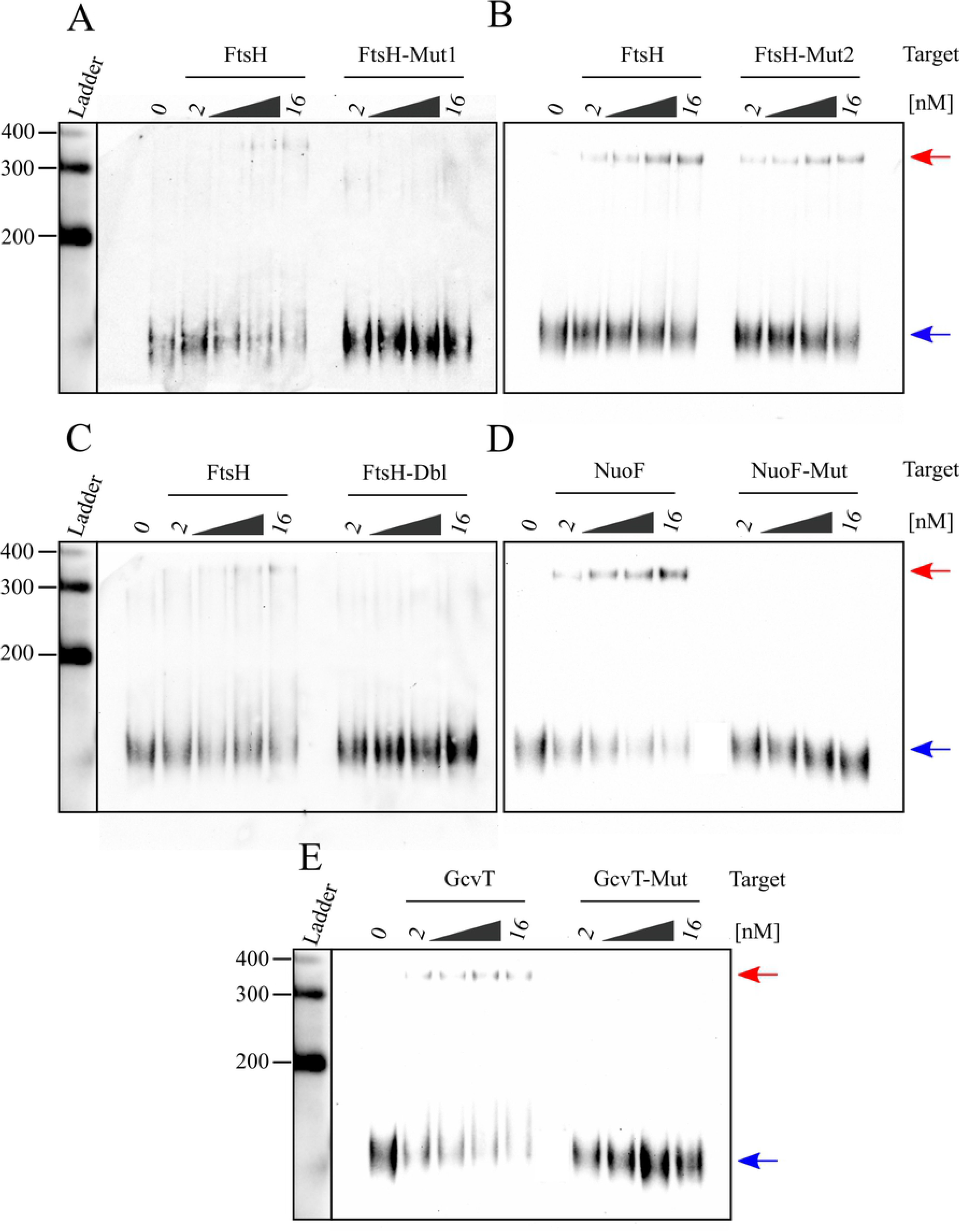
BbsR9 binds to ftsH, nuoF, and gcvT transcripts via specific GC-rich seed regions. RNA-RNA EMSAs showing dose-dependency of biotin-labeled BbsR9 binding to wild-type but not mutated, *in vitro-*transcribed segments of **A-C)** *ftsH*, **D)** *nuoF* and **E)** *gcvT*. Mutated regions correspond to those shown in **Fig 6B**, and “Dbl” specifies the double *ftsH* mutant. Red and blue arrows indicate bands corresponding to BbsR9 bound and unbound to target RNAs, respectively. All reactions contained 2 nM biotin-labeled BbsR9 in addition to increasing amounts of the indicated targets (0, 2, 4, 8, and 16 nM, respectively).

## Discussion

In this study, we performed an extensive transcriptomic analysis of *B. bacilliformis* grown *in vitro* then shifted to one of 10 distinct conditions that mimic environments encountered by the bacterium during its natural life cycle. We chose these conditions in order to control for a variety of environmental factors that may directly influence expression of certain sRNAs. For example, temperature (25°C, 30°C, 37°C), pH levels (pH 6, pH 7.4, pH 8.2), solid/liquid substrates, and presence of a blood-gas atmosphere (5% CO_2_, 2.5% O_2_, and 92.5% N_2_ at 100% humidity) were all examined. In addition, we included RNA-Seq experiments from experimental infections of low-passage human vascular endothelial cells (HUVEC) and fresh human blood samples (HB37 and HBBG). From these experiments, we discovered 160 sRNAs expressed by *B. bacilliformis* in at least one of the conditions tested.

Although we initially approached sRNA discovery using an automated approach, some clear-cut sRNAs were missed during the process. This issue led us to manually curate the 10 stranded RNA-Seq alignments, scanning each annotated gene, leader region, and IGR for aligned reads forming peaks that could represent novel sRNAs. These peaks were required to surpass a pre-determined read coverage threshold determined independently for each condition based on reads aligned to the *rpoD* (locus tag: BARBAKC583_RS04670) gene, which was consistently expressed across all 10 conditions (TPM ~300; **S4 Table**). We remained consistent by using *rpoD* as the housekeeping gene in qRT-PCR analyses (**S3 Fig**) and employing a 300 TPM threshold for the purpose of the UpSet plot (**Fig 1**).

The putative sRNAs identified were organized into three categories (IGR, *cis*-anti, or leader) depending on location of the corresponding sRNA locus. Each sRNA category has implications for its potential function. For example, IGR sRNAs are likely *trans*-acting with small seed regions that often bind multiple mRNAs. *Cis*-anti sRNAs most likely target the gene to which they are antisense, so target identification via algorithms such as IntaRNA would not be useful. Putative leader sRNAs are peaks that were identified sense to and in the 5’ UTRs of protein-coding genes. Although the identified peaks appeared distinct from those within the actual coding sequence, the possibility remains that these peaks are not *trans*-acting sRNAs. More likely, these leader RNAs may serve as *cis*-acting regulatory components, like riboswitches, which are co-transcribed with the downstream protein-coding gene and harbor regulatory stem-loops that influence translation of the respective transcript [55]. Determining whether the identified leader sRNAs are *cis*-and/or *trans*-acting elements would require further experiments such as Northern blots and 5’ RACE experiments to see if there is read-through into the downstream gene.

We performed Northern blot analyses on the putative leader sRNAs, BbspeF and BbsR7, and found that these are likely *cis-*acting leader RNAs, since the RNA sizes suggest read-through into the downstream gene (**Fig 2**). Northern blot analysis also verified the existence of six other sRNAs, although some of the results raise additional questions. For example, the presence of BbsR2 was detected, but the apparent band of ~450 bases is considerably larger than its predicted 284-base band (see **Fig 2**, **S3 Table**). Although we identified a putative promoter element for BbsR2, we did not identify a Rho-independent terminator, so it is possible that the sRNA extends further downstream than predicted. It was also unclear whether BbsR3-1 / BbsR3-2 represented two distinct sRNAs. However, Northern blot analysis utilizing a probe against BbsR3-1 confirmed that there was a single transcript produced whose length (~600 bp) was equal to the sum of the predicted sizes of BbsR3-1 and BbsR3-2, indicating that this locus probably produces a single sRNA species (**Fig 2**). The BbsR7 blot also requires explanation. Here, several bands were identified, including smaller bands of ~200 bases and a larger band of ~600 bases. Since BbsR7 is predicted to be a leader sRNA, it is possible that the smaller bands represent the sRNA being independently expressed, while the larger band may represent BbsR7 being co-transcribed with the downstream gene (BARBAKC583_RS01695), which is 225 bp long (**Fig 2**). We also probed in the BbsR11-1 / BbsR11-2 region to determine if the two corresponding RNA-Seq peaks represented two distinct sRNAs. In this case, the Northern blot showed a single band that corresponded only to the predicted size of BbsR11-1 (**Fig 2**), suggesting that the locus harbors two distinct sRNAs. Northern blots for BbgpI and BbsR9 produced single bands (**Fig 2**) that corresponded well to the estimated sizes of their respective peaks by RNA-Seq.

Among the sRNAs analyzed by Northern blot, Bbar45 and BbspeF are intriguing, non-coding RNA elements worthy of further characterization. Bbar45 belongs to the αr45 sRNA family first described in *Sinorhizobium meliloti*, but it is widely conserved in other Rhizobiales [56]. Functional characterization of sRNAs in the αr45 family has not been performed, although the *S. meliloti* αr45 can be co-immunoprecipitated with Hfq [57]. Since Hfq is an RNA chaperone that facilitates sRNA-mRNA interactions, we hypothesize that the *S. meliloti* αr45 sRNA may be *trans*-acting [58]. Here, we confirmed that Bbar45 is independently expressed from BbspeF, which lies immediately downstream (**Fig 2**). While this observation was previously observed in *S. meliloti*, it was unclear whether it was the case for other alphaproteobacteria [56]. Based on Northern blot results showing a transcript >700 bases (**Fig 2**), BbspeF is likely a leader RNA that is not independently expressed from its downstream gene. The *speF* leader RNA was initially discovered during a search for alphaproteobacterial riboswitches and was named for its upstream location relative to the *Bacillus subtilus speF* ortholog, which codes for an ornithine decarboxylase protein involved in polyamine biosynthesis [59]. However, metabolites of the polyamine biosynthesis pathway of *B. subtilus* were not shown to bind to the *speF* leader *in vitro*, leaving the element’s function unclear [59]. More experiments are needed to determine the regulatory role of the BbspeF leader as well as the function of the Bbar45 sRNA in *B. bacilliformis*.

An RNA secondary structure prediction of BbsR14 showed two stem-loops with nearly identical sequences of TTCCTCCTAA. Remarkably, these are anti-Shine-Dalgarno (anti-SD) motifs most often found in 16S rRNA, where they function in translational initiation. The presence of SD sequences outside of a ribosome binding site (RBS) is rare, as they are selected against in the context of mRNAs, since they can cause ribosome stalling due to hybridization with 16S rRNA [60, 61]. One way in which sRNAs regulate translation is to bind directly to the RBS to occlude the ribosome and inhibit translational initiation [31]. In most cases, this is accomplished via a seed region that overlaps the SD sequence and extends up and/or downstream [62]. The predicted BbsR14 secondary structure displays unique potential seed regions solely comprised of anti-SD sequences. We speculate that this arrangement could provide opportunities for indiscriminate translational repression by the BbsR14 sRNA.

We also analyzed each of the identified *B. bacilliformis* sRNAs and discovered that BB019, BB113, and BB125-2 possessed a single anti-SD sequence (CCTCCT). Interestingly, of the four sRNAs that contain anti-SD sequences, BbsR14 and BB113 were significantly upregulated at pH08 relative to pH06 (see **S3 Table**). Conditions of pH08 and pH06 were designed to simulate the initial and late stages of the sand fly after feeding, respectively. Thus, downregulation of translation may be advantageous for bacterial survival during initial stages within the sand fly’s midgut. As *B. bacilliformis* persists in the sand fly and infection proceeds, “gearing up” for a subsequent mammalian infection may occur as the insect prepares for another blood meal. Supporting this notion, we also identified 6S RNA as a sand fly-specific sRNA that was upregulated at pH08 vs. pH06, although not significantly (see **S4 Table**). 6S RNAs function by binding to and sequestering the RNA polymerase holoenzyme [63]. The resulting global repression of transcription during the initial stages of sand fly infection and, to a lesser extent, throughout a sand fly infection, could conceivably promote persistence of *B. bacilliformis* in the insect.

The mRNA target enrichment analyses for potential sand fly and infection-specific sRNAs provided insight into the regulation of pathways necessary for bacterial survival in these disparate environments. For example, targets of sand fly-specific sRNAs were significantly enriched for genes involved in the FAD-binding GO term and the biosynthesis of amino acids KEGG pathway (**Fig 3**). FAD-binding proteins include a wide array of proteins that participate in numerous biological processes. Enrichment of these genes may reflect a relatively low availability of FAD during residence in the sand fly. *B. bacilliformis* encodes a bifunctional riboflavin kinase/FAD synthetase (BARBAKC583_RS05700), and although this gene is relatively lowly expressed in all conditions tested, there is a downregulation of its expression under sand fly-like conditions (pH07, average TPM = 49.04) compared to human blood infections (HBBG, average TPM = 84.94). Enrichment of genes involved in the biosynthesis of amino acids is possibly explained by the likely downregulation of transcription and translation under sand fly-like conditions, where *B. bacilliformis* enters into a stationary phase that may promote persistence.

The human infection-specific sRNA targets were enriched in multiple GO terms associated with transferase activities, transporters, and the phospholipid biosynthetic process and KEGG pathways associated with glycerolipid/glycerophospholipid metabolism and nucleotide excision repair (**Fig 3**). Among these, there is a clear regulation of cell wall constituents during human infection conditions that would presumably be associated with morphological changes to the bacterium in the human host or perhaps as a means of expressing outer membrane proteins/transporters that aid in bacterial growth and replication during infection. This may very well also be in response to stressors encountered under these conditions, since nucleotide excision repair also seems to be significantly regulated by infection-specific sRNAs.

When analyzing mRNA targets of the infection-specific sRNAs, it was clear that numerous sRNAs were predicted to target the same mRNA in several cases (see **S5 Table**). For example, of the 19 presumed *trans*-acting, infection-specific sRNAs, three independently target BARBAKC583_RS04310 transcripts, coding for lysylphosphatidylglycerol synthetase; an enzyme previously shown to augment a pathogen’s defense against host cationic antimicrobial immune peptides [64]. Additionally, four of the 19 predicted *trans*-acting, infection-specific sRNAs target BARBAKC583_RS00395 transcripts, coding for cobaltochelatase subunit CobT, which is involved in the synthesis of cobalamin (vitamin B_12_), an essential coenzyme for many biological reactions [65]. It is difficult to ascribe roles to these mechanisms without knowing whether the sRNA-mediated regulation is positive or negative, although it is worth noting that redundant targeting is not a result of sRNA duplication, and each predicted binding site on these transcripts is unique. We hypothesize that redundant regulation of particular mRNAs may serve to “hyper-regulate” protein production in response to subtle differences in environmental cues. This kind of redundant regulation of mRNAs from multiple “sibling sRNAs” has been described in other pathogens, so further research into the function of sibling sRNAs of *B. bacilliformis* could be fruitful [66].

We found conservation of some sRNAs among other alphaproteobacteria species using discontinuous megaBLAST analysis (**S3 Table**). Unfortunately, we were only able to analyze IGR and leader sRNAs, since *cis*-anti sRNAs showed broad sequence conservation due to their close linkage to protein-coding genes. The majority of analyzed sRNAs was unique to *B. bacilliformis*, while the BB036 sRNA group was unique to the KC583 strain of *B. bacilliformis*. Five other sRNAs were widespread in *Bartonella* spp., including BbsR9 which was characterized in this study. Conservation of BbsR9 in other *Bartonella* spp. further highlights its potential importance. Since *Bartonella* spp. are typically transmitted to mammals by various arthropods (ticks, sand flies, fleas, lice, etc.), it is possible that BbsR9 plays a role in persistence in many vectors. Seven more sRNAs were found in additional alphaproteobacteria, including ubiquitous sRNAs like 6S RNA and tmRNA, conserved alphaproteobacteria sRNAs like Bbar45, and the tRNA_Arg_^CCU^ group I intron.

BbgpI is a member of a tRNA_CCU_^Arg^ group I intron family first identified in *Agrobacterium tumefaciens* and later found in other alphaproteobacteria [67, 49]. Group I introns are selfish genetic elements that insert into tRNAs, rRNAs, and protein-coding genes. Although group I introns are ribozymes and RNA splicing is auto-catalytic, they sometimes require protein co-factors for self-splicing *in vitro*, and it is presumed that all group I introns require protein co-factors to some extent for splicing *in vivo* [50]. Here, we have demonstrated that BbgpI self-splices *in vitro* and is spliced *in vivo*. Furthermore, we have shown that BbgpI is not located in an IGR as presumed, but rather within an unannotated tRNA_CCU_^Arg^ gene. Since the flanking tRNA_CCU_^Arg^ gene retains all necessary tRNA domains (see **S4C Fig**), we predict that the tRNA is functional following intron splicing. This novel tRNA gene might have implications for future analyses of *B. bacilliformis* involving codon bias, conservation of tRNA genes, amino acid scavenging, etc. Furthermore, this discovery suggests further optimization may be required for current tRNA scanning algorithms.

We also characterized the targeting and molecular interactions of BbsR9, as the sRNA was only appreciably expressed under pH06, pH07, and pH08 conditions (**S4 Table**). For reference, these conditions reflect a liquid blood / serum environment at 30°C (**Table 1**) and simulate the sand fly’s midgut following a blood meal. It is interesting to note that the Pl30 condition is identical to pH07 except that Pl30 represents a solid medium. Furthermore, Northern blot analyses indicated that, in addition to the liquid medium requirements, BbsR9 expression was restricted to temperatures < 37°C (**S5B Fig**). The regulatory mechanisms that facilitate such an expression pattern warrant further investigation.

We verified several mRNA targets of BbsR9 using RNA-RNA EMSAs. Among the targets were transcripts of the *ftsH*, *nuoF*, and *gcvT* genes. First, *ftsH* codes for the FtsH zinc metalloprotease; a membrane-anchored, universal protease with various functions. FtsH has been extensively studied in *E. coli*, where it is the only protease essential for survival [reviewed in 68]. FtsH has also been described as required for regulation of optimal ratios of phospholipids and lipopolysaccharides in the outer membrane [68]. Whether BbsR9 regulation of *ftsH* transcripts is involved in bacterial protein turnover and/or modulation of membrane architecture in the context of the sand fly is unknown, but would be interesting to investigate. Second, the *nuoF* gene codes for the NADH-quinone oxidoreductase subunit F, a component of the type I NADH dehydrogenase enzyme and the initial step in the electron transport chain. NuoF is a component of the peripheral fragment of the NADH dehydrogenase complex and plays a role in oxidation of NADH to generate a proton motive force [reviewed in 69]. Regulation of *nuoF* transcripts could conceivably play a role in helping to establish the stationary phase as *B. bacilliformis* persists in the sand fly. Finally, we determined that BbsR9 targets transcripts of *gcvT*, which codes for the glycine cleavage system aminomethyltransferase, GcvT. The glycine cleavage system responds to high concentrations of glycine, breaking the amino acid down to CO_2_, ammonia, and NADH [70]. In addition to redox reactions, NADH can be used to produce energy through cellular respiration. Of note is the potential interplay between sRNA targeting of *nuoF* and *gcvT* transcripts in this regard. Interestingly, the glycine cleavage system has been implicated in contributing to bacterial persistence in animal and plant hosts [71]. In fact, *gcvT* is essential for persistence of a closely-related pathogen, *Brucella abortus*, in its animal host [72]. However, to our knowledge, the role of a glycine cleavage system in pathogen persistence in its arthropod vector has not been explored, to date. It is conceivable that *B. bacilliformis* utilizes regulation of *nuoF* and *gcvT* to fine-tune levels of NAD+/NADH, thereby contributing to regulation of metabolism and persistence of the bacterium in the sand fly.

This study has provided further insight into the regulation of numerous processes by *B. bacilliformis* in response to conditions encountered in the context of its sand fly vector and human host. We believe the results provide a strong foundation for future studies examining sRNA-mediated regulation in *B. bacilliformis* and the regulatory mechanisms required for vector-host transmission.

## Acknowledgments

The authors wish to thank Patty Langasek and Auguste Dutcher for technical assistance. This work was supported by NIH grant R21AI128575 (to MFM). RR was supported by NIH grants AI133023 and DE028409.

## Supporting information

**S1 Fig. *B. bacilliformis* RNA-Seq PCA plot.** Axes indicate the percentage of total variance that can be accounted for by two principle components. Colored dots indicate the retained biological replicates of the RNA-Seq analyses, and their distance apart is representative of overall relatedness in gene expression profiles. Experimental conditions are shown on the right.

**S2 Fig. *B. bacilliformis* sRNAs group into specific expression patterns.** Heatmap of *B. bacilliformis* sRNA TPMs across the tested conditions (shown at the bottom). sRNAs group vertically based on similarity in expression patterns. Conditions group horizontally based on similarity in overall expression patterns. The log_10_ of the TPM value for each sRNA is indicated by a color gradient.

**S3 Fig. qRT-PCR confirmation of differential expression of several identified sRNAs.** Faceted bar graph displaying the number of sRNA transcripts / *rpoD* transcript for select, differentially-expressed sRNAs and BB024, which was not shown to be differentially expressed. The condition / source of the total RNA is noted on the x-axis. Significance was determined by students t-test (N = 9; * = p<0.05, ** = p<0.01, *** = p<0.001).

**S4 Fig. BbgpI is a group I intron inserted into an unannotated tRNA_CCU_^Arg^ gene of *B. bacilliformis*. A)** Nucleotide sequence of BbgpI (bolded and underlined) and flanking chromosomal regions. Primer binding sites used for *in vitro* transcription (IVT) and PCR assays designed to show splicing of BbgpI *in vitro* and *in vivo* are indicated. **B)** Sequence of BbgpI outlining the conserved, characteristic stem structures (P1 to P9) with putative base pairings highlighted in green and yellow. Nucleotides predicted to participate in base pairing are bolded and underlined. **C)** Sequence coding for the tRNA_CCU_Arg immediately flanking BbgpI. The two bolded underlined nucleotides represent the ends of the spliced out BbgpI. Conserved tRNA features are also outlined.

**S5 Fig. BbsR9 is a sand fly-specific sRNA. A)** Nucleotide sequence of the *bbsR9* gene with predicted promoter elements and Rho-independent terminator plus experimentally-determined TSS’s, highlighted in various colors or underlined, respectively. An asterisk indicates the alternative TSS found by 5’ RACE analysis. **B)** Northern blot analysis of BbsR9 expression under the indicated conditions. The RNA ladder (2 min exposure) and resolved total RNA samples (30s exposure) were from the same blot but imaged using different exposure times.

**S1 Table. Bacterial strains, primers, and plasmids used in the study.**

**S2 Table. Quality control results for *B. bacilliformis* RNA-Seq analyses.**

**S3 Table. Putative sRNAs identified in *B. bacilliformis* by RNA-Seq analyses.**

**S4 Table. Average TPMs of identified *B. bacilliformis* sRNAs.**

**S5 Table. Predicted IntaRNA targets of *B. bacilliformis* infection-specific sRNAs.** An “X” indicates transcripts of the indicated gene to which the sRNA is predicted to bind (p < 0.01). Targets with a FDR < 0.05 are indicated with a red “X”.

**S6 Table. Predicted IntaRNA targets of *B. bacilliformis* sand fly-specific sRNAs.** An “X” indicates transcripts of the indicated gene to which the sRNA is predicted to bind (p < 0.01). Targets with a FDR < 0.05 are indicated with a red “X”.

